# More Severe Loss-of-Function Sodium Channel induced by Compound SCN5A R965C and R1309H Mutants lead to Complex Familial Arrhythmia Syndrome

**DOI:** 10.1101/725945

**Authors:** Yubi Lin, Jiading Qin, Yuhui Shen, Bin Jiang, Xiufang Lin

## Abstract

**Objective:** A complex familial arrhythmia syndrome was identified in a Chinese Han family, characterized by sick sinus syndrome, progressive conduction block, atrial fibrillation, atrial standstill and Brugada syndrome. The underlying mechanism associated with the genetic mutation was investigated.

**Methods:** Genetic analysis was conducted for probands in the coding and splicing regions of genes susceptible to cardiac arrhythmia diseases. The stable cell lines overexpressing wild-type (WT) or mutant SCN5A gene were generated in HEK293T cells and electrophysiological recordings were performed to evaluate the functional changes of mutant Na_V_1.5 channels.

**Results:** We identified a rare double R965C and R1309H mutations in Na_V_1.5 channel, as a heterozygous and compound pattern, locating in the same chromosome. Compared to WT, a single mutation of R965C or R1309H, the compound mutations of R965C-R1309H most dramatically reduced the peak current of Na_V_1.5 channel with testing potentials ranging from −35 mV to 25 mV. Notably, the maximal peak current of Na_V_1.5 channel carrying R1309H only and R965C-R1309H mutations displayed significant decreases by 31.5% and 73.34%, respectively, compared to WT. Additionally, compared to single mutations of R965C or R1309H, the R965C-R1309H mutations resulted in more obviously depolarization-shifted activation and hyperpolarization-shifted inactivation, also caused most significantly changed time constant, V_1/2_ and slope factor of activation and inactivation.

**Conclusions:** The compound mutations of R965C-R1309H led to more dramatically reduced current density of Na_v_1.5 channel, as well as more depolarization-shifted activation and hyperpolarization-shifted inactivation than single mutation of R965C or R1309H, which may provide more understanding on the molecular mechanisms underlying the familial arrhythmia syndrome.

**Author summary:** Several limited types of research about the interaction of compound mutations and the mechanism of interaction remains controversial. Here we report a novel compound mutation R965C-R1309H in SCN5A gene.The compound mutations of R965C-R1309H markedly reduced the Na^+^ current density of Na_v_1.5 channel and caused significant depolarization-shifted activation and hyperpolarization-shifted inactivation, suggesting that R965C mutation aggravated the pore gating changes of sodium channel containing R1309H mutation. The R965C located in the S4-like region in the intracellular loop between DII and DIII, and the R1309H located in the S4 helix of DIII as voltage sensor, which thus potentially had synergistic and interacting effects on pore gating properties of the sodium channel. Additionally, the complex familial arrhythmia syndrome simultaneously complicated of early young onset of arrhythmia and harmful prognosis of recurrent ischemic cerebral stroke, even though effective Warfarin anticoagulation.

## Introduction

The SCN5A gene encoding the pore-forming α-subunit of the cardiac voltage-gated sodium channel (Na_v_1.5) is responsible for the generation and rapid propagation of action potentials as a rapid inward Na^+^ current (I_Na_), in the heart(1). This highly conserved 100 kb gene is located on chromosome 3 and contains 28 exons encoding four homeodomains, including six transmembrane segments (S1–S6) forming a channel pore that determines the selectivity and permeability of the sodium channel(2). As reported in the Databases of Online Mendelian Inheritance in Man (OMIM), pathogenic mutants of SCN5A encoding the cardiac sodium channel have been linked to a wide range of inherited and malignant arrhythmia syndromes, such as type 3 long QT syndrome(3), Brugada syndrome(4, 5), progressive conduction disease(6), sick sinus syndrome, atrial fibrillation, atrial flutter, atrial standstill, ventricular fibrillation, dilated cardiomyopathy, right ventricular cardiomyopathy(7, 8) and sudden infant death syndrome(9). Mixed phenotypes are frequently referred to as cardiac sodium channel overlap syndrome, heritable/familial arrhythmia syndrome or SCN5A syndrome induced by a single or several SCN5A mutations.

Here, we report a Han Chinese family presenting with complex familial arrhythmia syndrome, characterized as sick sinus syndrome, progressive conduction disease, atrial fibrillation, atrial flutter, atrial standstill and Brugada syndrome, and investigate the genetic background and underlying pathogenic mechanism associated with SCN5A mutation.

## Results

### Clinical presentation

The familial pedigree was shown in **Figure 1**. Proband III: 7, 10-year-old boy, suffered from sick sinus syndrome (SSS) and high-degree atrioventricular block (AVB) with syncope and dizziness (**Figure2A, B**), thus permanent cardiac pacemaker was implanted. One year later, atrial fibrillation was developed (**Figure 2C**) and recurrent even after catheter ablation. His father II:3 complicated of atrial flutter with fasting ventricular rate (**Figure 2D**) at the age of 41, which was treated by catheter ablation. He experienced SSS and intermittent third-degree AVB (**Figure 2E**) during one-year follow-up and ultimately was implanted with a permanent cardiac pacemaker. He experienced three episodes of ischemic cerebral stroke due to persistent atrial fibrillation and atrial standstill(**Figure 2F**), which led to the disorder of limb activity, despite orally effective anticoagulation of Warfarin. Recent ECGs of II: 3 appeared with ventricular pacing dependency (**Figure 2D**), suggesting progressive and complete AVB of the third degree. ECGs of II:2 (**Figure 2F**) showed type I and II Brugada pattern (**Figure 2G**). II:2 had a history of syncope associated with overexertion and lack of sleep without ECG recordings, but refused electrophysiological examination and further Holter recording. Although III:3 had no malignant event occurred, he was demonstrated with sinus bradycardia in the ECG of daytime (**Figure H, I**). Moreover, the mean heart rate of III:3 was ≤40 beats/min in the daytime of 24-hour Holter recording. III:4 and I:1 had an asymptomatically first degree/intermittent second degree, and first-degree AVB, respectively, in 24-hour Holter recording. The ECGs and 24-hour Holter recording of other members were normal. There were no significant abnormalities of cardiac structure, evaluated by echocardiographic imaging in this family.

**Figure 1:**
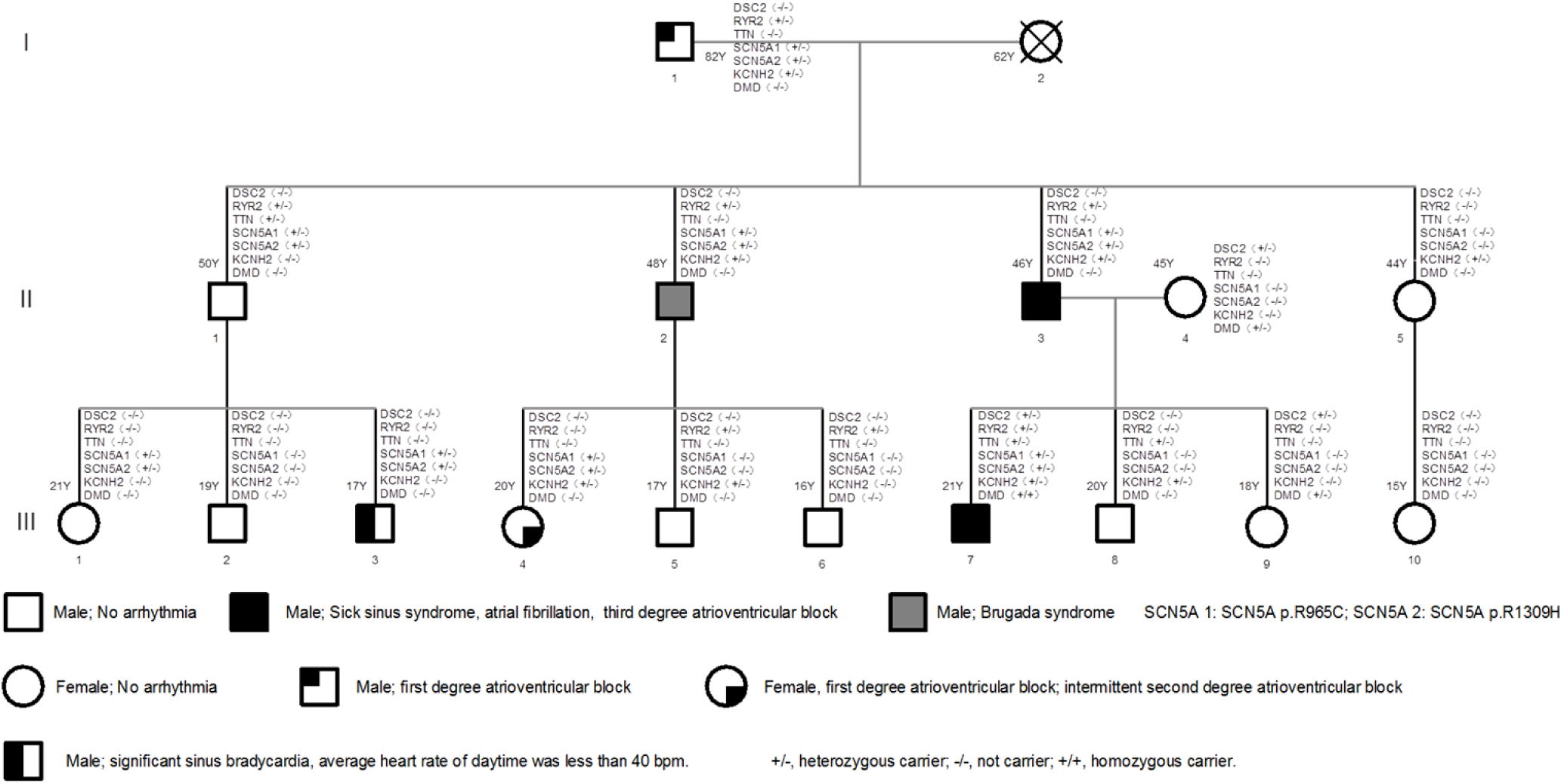
The familial pedigrees with SCN5A mutations. The gene abbreviations were as follows: DSC2, desmocollin 2; RYR2, ryanodine receptor 2; TTN, titin; SCN5A, sodium voltage-gated channel alpha subunit 5; KCNH2, potassium voltage-gated channel subfamily H member 2; DMD, dystrophin. ±, the heterozygous carrier with mutations; -, no carrier with mutations.

**Figure 2.**
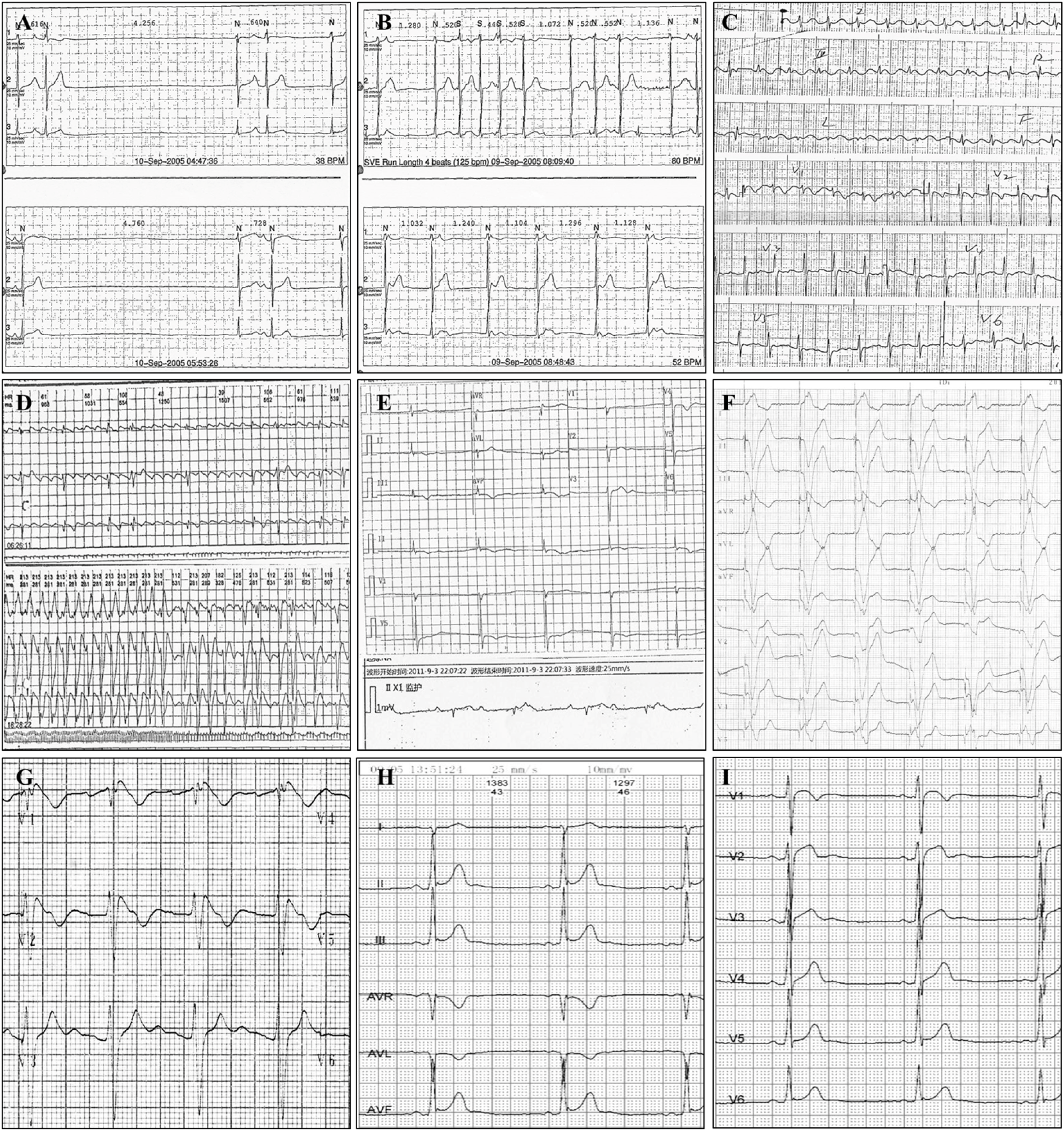
The electrocardiograms of probands. The ECGs of III:7 and II:3 were showed in A-C and D-F, respectively. A) sinus arrest for approximately 4.25 (top) and 4.76 (bottom) seconds, junctional escape beat and premature atrial beat; B), top, sinus arrest, junctional escape beats, frequent premature atrial beats and short escapes of atrial tachycardia; button, sinus arrest, and junctional escape rhythm. C) atrial fibrillation with a fast ventricular rate and intraventricular block. D) top, atrial flutter with 2:1 to 6:1 atrioventricular conduction; button, atrial flutter with fast ventricular rate. E) top, sinus arrest, and junctional escape rhythm; button, third-degree atrioventricular block, and junctional escape rhythm. F, atrial standstill, persistent ventricular pacing, termed as ventricular pacing dependence. G) ECG of II:2, Brugada-like ECG of type I. H) ECG of III:3, sinus bradycardia during the daytime.

### Genetic background

After filtering single nucleotide polymorphisms (SNPs) with minor allele frequencies ≥0.001, III:7 was identified with heterozygous mutants of SCN5A (c.2893C>T, R965C), SCN5A (c.3926G>A, R1309H), RYR2 (c.5614G>A, D1872N), KCNH2 (c.3094C>T, R1032W), DSC2 (exon15: c.C2497T:p.R833C), RYR2 (exon37: c.G5614A:p.D1872N), TTN (exon2: c.G13A:p.A5T) and DMD (exon9: c.G1140C:p.K380N), as shown in **Table 1**. The RYR2, SCN5A, and KCNH2 genes were associated with hereditary arrhythmia and cardiac ionic channelopathies, while the DSC2, DMD, and TTN genes led to susceptibility to cardiomyopathies(14, 15). Polymerase chain reaction-based sequence of SCN5A showed C-T transversion at codon 2893 of exon 17 and G-A transversion at codon 1309 of exon 22, resulting in the substitution of Arg for Cys (R965C) and Arg for His (R1309H). The minor allele frequencies of SCN5A R965C and R1309H were not obtained (0%) in Frequency of 1000 Genomes database. The risk of pathogenesis was predicted as “Damaging” by the polyphen and SIFT algorithms. All of these mutations mentioned above were validated by Sanger sequencing in all families. The genotype-phenotype coseparation analysis revealed that, since the two heterozygous mutations of R965C and R1309H (compound mutation of R965C-R1309H) passing down the generations, as a heterozygous and compound pattern, suggesting that the two mutations located in the same chromosome. The members of this family, including I:1, II:1-3, III:1, III:3, III:4 and III:7, carried the compound mutations. II:5 carrying with KCNH2 but not SCN5A mutations had normal ECGs and cardiac structure without arrhythmia at the age of 44. III:3 carrying with the compound mutations of SCN5A but not KCNH2 and RYR2 mutations complicated of early sinus node dysfunction in the daytime, suggesting KCNH2 and RYR2 were not the necessary factors of pathogenesis. Furthermore, the mutations of DSC2 and DMD were also validated in II:4, who had no kinship related to this family. In addition, the mutation of TTN encoding the cardiac structure protein of titin was naturally common in the population. Also, there was no clinical evidence of cardiomyopathies for this family. R965C was located within the intracellular loop between domains II and III (DII-III) of Na_v_1.5 protein encoded by SCN5A gene. R1309 was localized in the S4 helix of the repeat III transmembrane region (DIII) and function as a voltage sensor in the sodium channel. The conservation analyses demonstrated that R965 and R1309 were highly conserved across all species.

### Electrophysiological characteristics

To detect the effect of R965C-R1309H mutations on Na_V_1.5 channel function, WT Na_V_1.5 channel and mutant Na_V_1.5 channels carrying R965C (Na_V_1.5_R965C_), R1309H (Na_V_1.5 _R1309H_) or R965C-R1309H (Na_V_1.5_R965C-R1309H_) were stably transfected in HEK293T cells, respectively, followed by whole-cell patch clamp recordings. Representative traces of the four groups were shown in **Figure 4A**.

**Figure 3.**
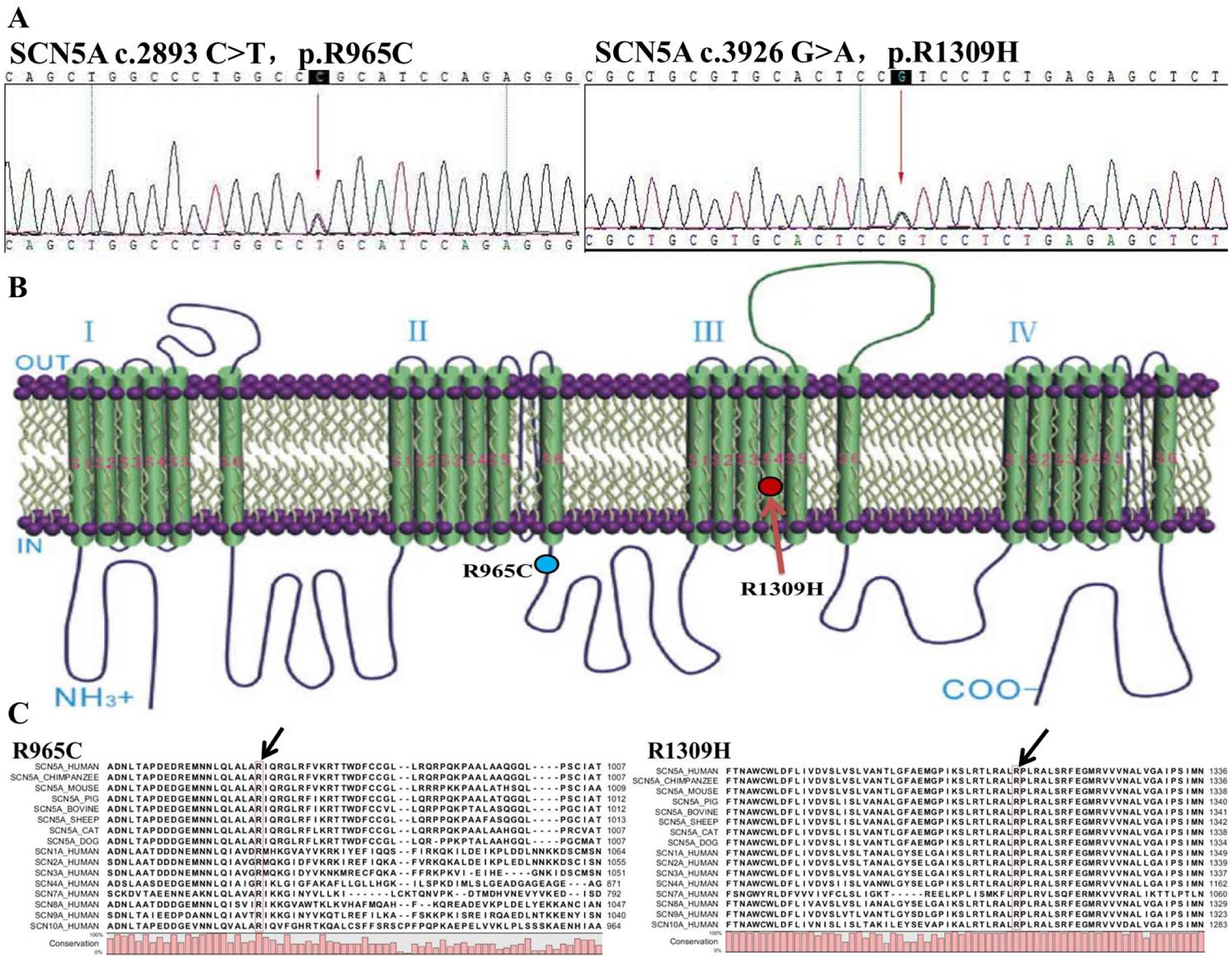
The molecular genetics of SCN5A linked to familial arrhythmia syndrome. A) Sanger sequencing for both of R965C and R1309H mutations. B) R965C located in the cytoplasmic loop between domain II and III. R1309H located in S4 the helix of domain III as a voltage sensor. C) Arginine in 965 and 1309 sites of an amino acid sequence of Na_v_1.5 protein was conserved in all species.

**Figure 4.**
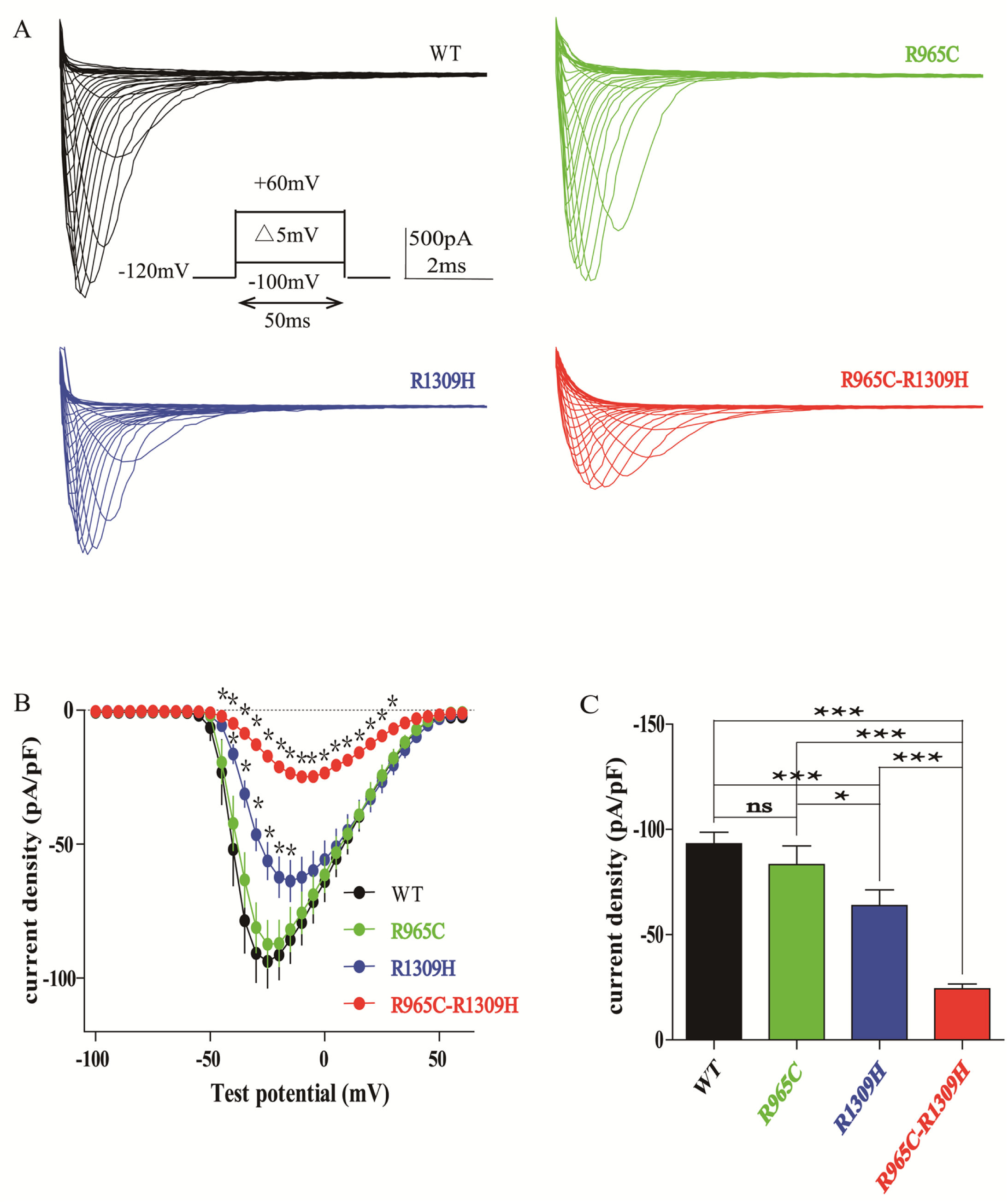
The peak currents of Na_v_1.5 channels recorded in HEK293T cells. Voltage clamping of wild-type and mutant Nav1.5 channels-transfected HEK293T cells at 32°C. A) Representative traces of WT and mutant Nav1.5 sodium channel. B) Current density and voltage (I-V) relationship. C) The peak current of WT and mutant Nav1.5 sodium channel. *p*-value less than 0.05 was considered to be statistically significant. Compared to the WT group, * *p*<0.05; ** *p*<0.01; *** *p*<0.001.

Consistent with previous literatures reported(16, 17), as shown in the Figure **4A-C and Table 2**, the R1309H mutation, but not R965C mutation, significantly reduced the Na^+^ current density at a series of test potentials ranging from −45 mV to −15 mV when compared to the WT group (**Supplement 1**). More importantly, we found that R965C-R1309H mutations resulted in a more severe reduction in Na^+^ current density ranging from −45 mV to 30 mV compared to the WT group. Even compared to R965C or R1309H mutation, the R965C-R1309H mutations still had a significantly lower Na^+^ current density ranging from −35 mV to 25 mV. In details, the peak current of Na_V_1.5 channel with R1309H and R965C-R1309H mutations were dramatically reduced by 31.5% (−63.77±7.59 pA, P<0.001) and 73.34% (−24.83±2.32 pA, P<0.001), respectively, while the peak current of Nav1.5 channel with R965C remain unchanged (−87.37±8.83 pA), compared to the WT group (−93.14±5.55 pA) (**Figure 4C**).

In addition, the R965C-R1309H mutations also caused a significant shift in the steady-state activation of Na_v_1.5 channel (**Supplement 2**). The R1309H mutation caused a right-shift effect on the activation curve **(Figure 5A**), depolarized the V_1/2_ and enlarged the slope factor (**Table 3**). Furthermore, the time constant of R1309H mutation was also amplified ranged from −45mV to −30mV when compared to WT group, which indicated that Na_v_1.5_R1309H_ channel was activated at the more depolarized membrane potential and took a prolonged time to run up to the peak current. However, R965C mutation did not induce significant changes among all the above aspects. Those results were basically consistent with previous studies(16, 17). As for R965C-R1309H mutations, compared to R1309H mutation, it induced more significant depolarization shift of activation curve, V_1/2_ and slope factor, suggesting that Na_v_1.5_R965C-R1309H_ channel was activated at the much more depolarized membrane potential and displayed slower activation (**Supplement 3**). However, the time constant of Na_v_1.5_R965C-R1309H_ channel was significantly amplified compared to WT group between −40mV and −30mV (**Figure 5B**) but shortened at the test potential of −45mV and −40mV compared to Na_v_1.5_R1309H_ channel. We hypothesized that this may be due to the small current of Na_v_1.5_R965C-R1309H_ channel that led to a decreased time to reach the peak current.

**Figure 5.**
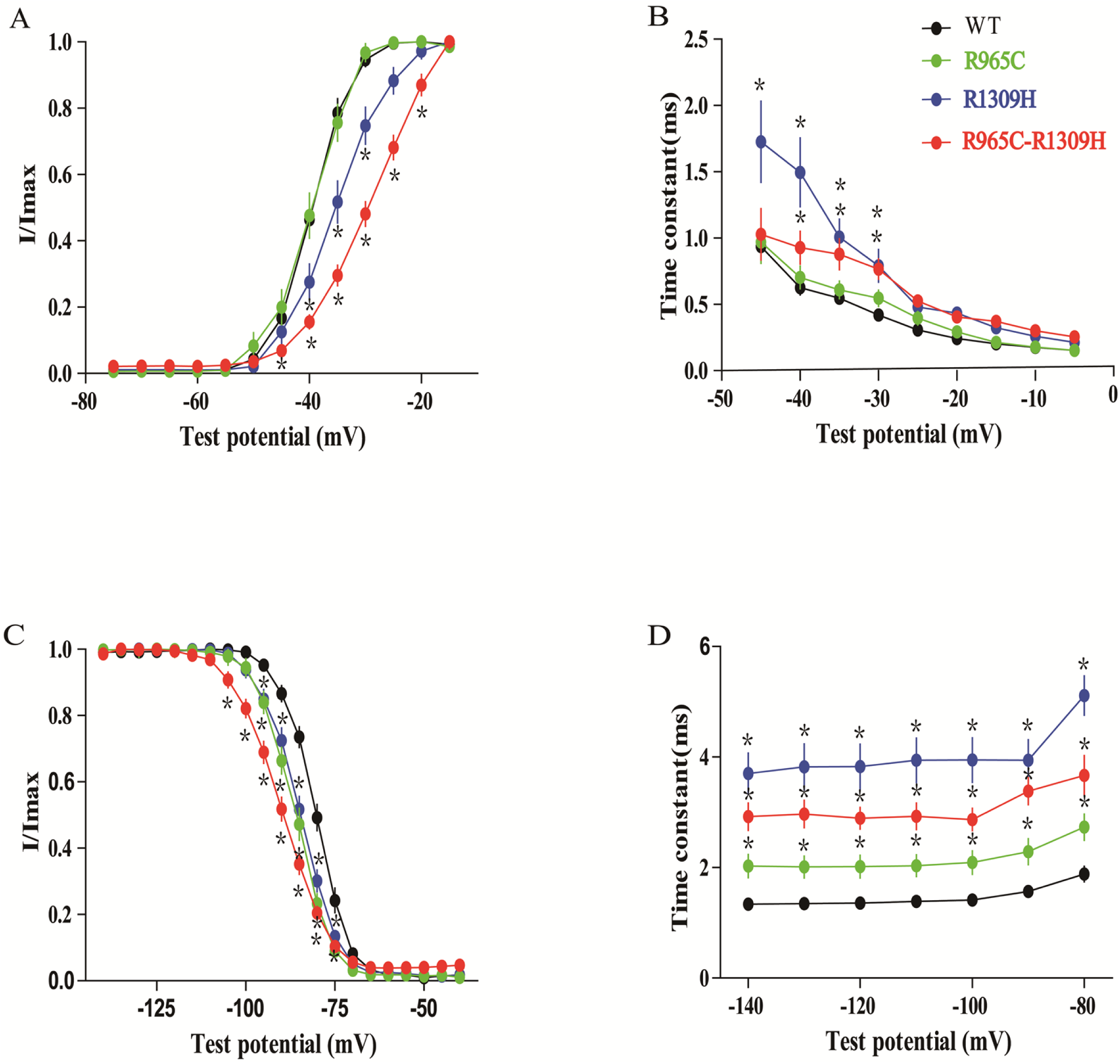
Kinetics of sodium channel in the steady-state activation and inactivation. A) Steady-state activation as a function of voltage. B) Relationship between tau (τ) and test potential for activation. C) Steady-state inactivation as a function of voltage. D) Relationship between tau (τ) and test potential for steady-state inactivation. Compared to the WT group, and *p*-value less than 0.05 was considered to be statistically significant (*).

The R965C-R1309H mutations also caused more intense damage in the steady-state inactivation of the sodium channel (**Supplement 4**). Both of R965C and R1309H mutations showed most left-shift inactivation curve and hyperpolarized inactivation V_1/2_ respectively, which suggested a reduced availability of open channel. The time constants of R965C and R1309H mutations were significantly prolonged at the test potential between-140mV and −80mV, respectively, which mean the peak amplitude took longer time to decay (**Supplement 5**). And the slope factor (K) of R1309H mutation was significant amplified but not of R965C mutation when compared to the WT group. These above data were keeping with the previous studies(16, 17). We were surprised to find that the inactivation curve of R965C-R1309H mutations most significantly shifted to the left with the increase of V_1/2_ hyperpolarization and K value (**Figure 5C, Table 3**). Compared with the other two single mutations, the inactivation curve of the R965C-R1309H mutations also shifted further to the left, accompanied by further hyperpolarization of V_1/2_ and a further increase of K value, which suggested a much lower availability of open channel. The time constant of R965C-R1309H mutations was significantly longer than that of the WT group and was between R1309H and R965C mutations (**Figure 5D**). As for why the time constant of the R965C-R1309H mutations was not larger than that of both single mutations, we think it was related to the lowest peak current of the R965C-R1309H mutations.

### Intracellular trafficking analysis by confocal imaging

The construct used in this study had a flag epitope fused to the C-terminus of Na_v_1.5 protein so that we might regard antibody that recognized flag as the one that targets Na_v_1.5 protein. The construct also generated GFP protein respectively and we might, therefore, regard the presence of GFP protein as the presence of transfected cells. Similar intracellular distribution for WT and mutant SCN5A proteins and the presence of red fluorescence at the periphery of cells were observed. This suggested that a single mutation of R965C or R1309H, as well as the compound mutations of R965C-R1309H, were trafficking competent (**Figure 6**). There were some red spots in the cell which might be the immature proteins in ER pool.

**Figure 6.**
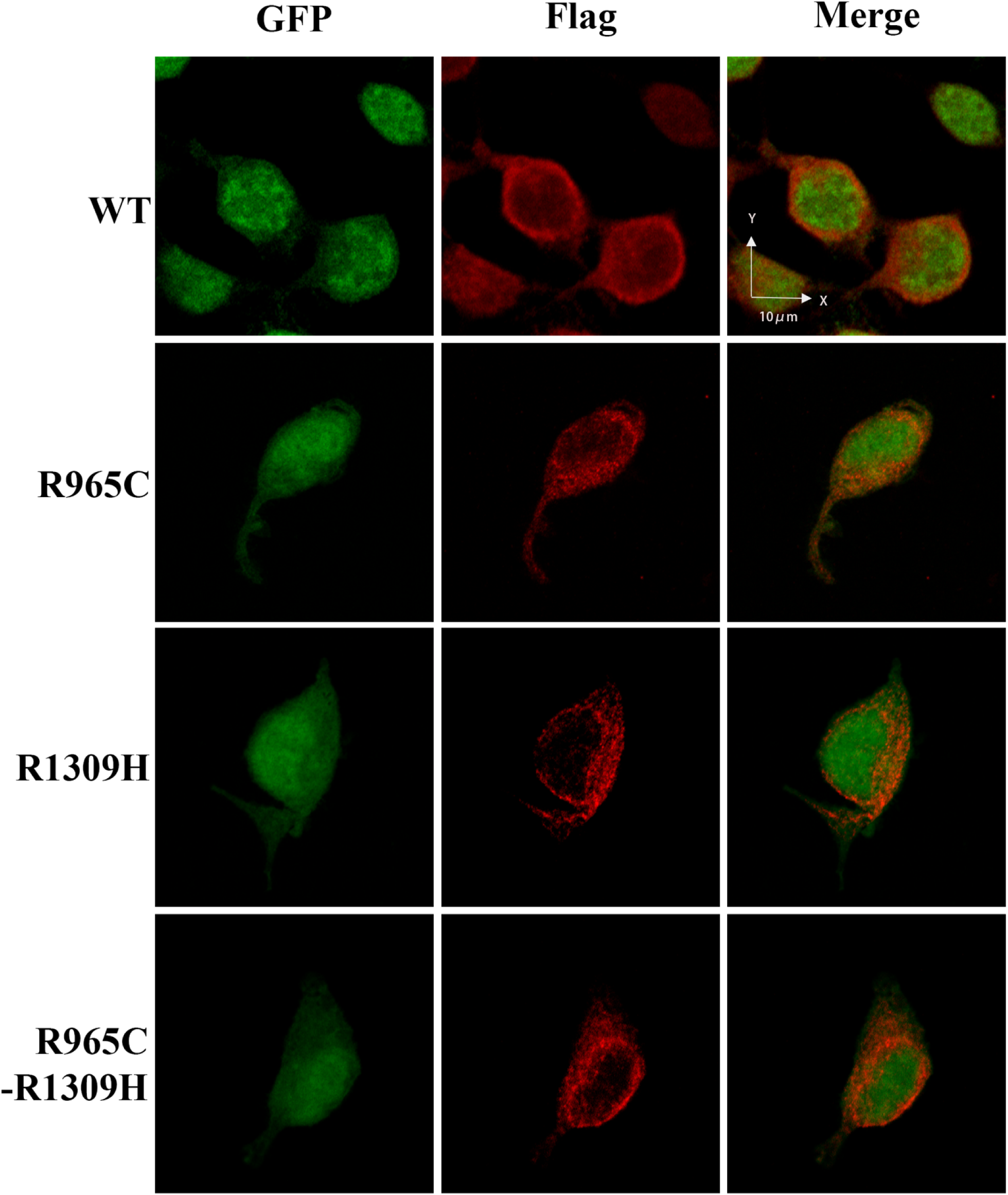
Immunocytochemistry staining revealed by confocal image. The total cells were analyzed for WT, R965C, R1309H and the compound mutations of R965C-R1309H respectively. The presence of red fluorescence (anti-flag antibody) at the cytoplasmic membrane indicated that Nav1.5 protein of WT, as well as other mutations, were all trafficking competent and locating to the membrane. The red spot at the center of the cell might be immature Nav1.5 protein in the endoplasmic reticulum.

## Discussion

While the R965C and R1309H mutations of SCN5A gene have been individually reported, we described for the first time the clinical presentation, biophysical properties and the interaction of the heterozygous and compound mutations of R965C-R1309H in the same chromosome, which more significantly reduced current density of I_Na_, led to more depolarized activation and hyperpolarized inactivation in a synergistic way. These changing pore gating properties would explain the reason for the severe clinical phenotypes of complex familial arrhythmia syndrome, characterized as sick sinus syndrome, atrial fibrillation, progressive conduction disease, atrial standstill and Brugada syndrome. The carriers individually exhibited one or multiple phenotypes due to the difference in penetrance of both mutations. Notably, the carriers complicating of atrial fibrillation and atrial standstill had apparently increased the risk of recurrent cerebral embolisms.

### Compound mutations of R965C and R1309H synergistically aggravated pore gating properties

According to previous studies, although R965C mutation carried by a sporadic case of Brugada syndrome didn’t significantly change the current density of I_Na_, it produced sodium channel with impaired steady-state inactivation, slower inactivation, and recovery from inactivation. The substitution of R965C from the positively charged arginine to a neutral cysteine might influence channel gating properties(17). In another family, the proband carrying homozygous R1309H mutation which inherited from the parents of close cousins, showed with complex clinical phenotypes characterized as follows: one proband had phenotypes of atrial tachycardia, ventricular tachycardia, Brugada syndrome-like ST segment and T wave, and first degree atrioventricular block; the second proband was demonstrated with sinus bradycardia, P wave prolongation, first degree atrioventricular block and QRS prolongation, also suffered from atrial septal defect and ventricular septal defect requiring surgical closure; another sibling of 2-month old experienced sudden cardiac death induced by diarrheal illness was speculated to carry with homozygous R1309H mutation, although the genotype was unknown. The R1309H mutation significantly reduced the current density of I_Na_, hyperpolarized inactivation, slowed recovery from inactivation, and increased the I_NaL._ It was noted that the carriers with heterozygous R1309H mutation showed slightly abnormal ECGs without arrhythmia events(16). In our study, we interestingly found that, the probands of II:3 and III:7 carried the compound mutations of R965C-R1309H locating in the same chromosome (while the other homologous chromosome was normal without SCN5A mutation), serve as heterozygous and compound pattern, and occurred with sick sinus syndrome, progressive conduction disease, atrial fibrillation, atrial standstill, whose phenotypes different from that of heterozygous R965C or R1309H mutation. Based on these phenomenons of obvious phenotype difference and severity of illness, we speculated that R965C and R1309H may have a collaborating effect on sodium channel with each other in the probands of this family. In order to further explore the synergistic effect for R965C and R1309H, we constructed stable cell lines of HEK293T transfected with viral vectors carrying with R965C, R1309H, and R965C-R1309H mutations, which therefore avoided the imprecise influence on sodium current due to different and low transfecting efficiency of transient transfection by ordinary plasmids. As we expected, compared to the WT group, we got similar results consistent with the two previous studies for R965C or R1309H mutation alone. Interestingly, no matter whether comparing to WT, R1309H or R965C mutation, the compound mutations of R965C-R1309H led to the most malignant loss of function of the sodium channel, including the lowest peak current density of I_Na_, slowest and depolarized activation, and most hyperpolarized inactivation, suggesting most apparently reduced availability of open sodium channel. Especially, the maximal amplitudes of I_Na_ of R1309H and R965C-R1309H mutations reduced to 31.5% and 73.34% of the WT group. The excessive reduction of I_Na_, depolarized activation, hyperpolarized inactivation would obviously decrease the action potential upstroke velocity and conduction velocity, thereby lead to cardiac conduction diseases including incomplete dysfunction or even completed loss function of the sinus node, atrial propagation, atrioventricular node, and intraventricular propagation. The changes of pore gating properties in our study had been similar to these previous studies(18-21), and consisting of the genotype-phenotype correlations in this family. In this family, not only R965C mutation but also R1309H and R965C-R1309H mutations could lead to Brugada syndrome or Brugada syndrome-like ECG. These three kinds of mutations characteristic of incomplete or completed loss of function might subsequently produce transmural gradients of membrane potentials of the right ventricle during early repolarization caused by subepicardial phase 0 block and discontinuous transmural conduction. At the same time, the late slow upstroke action potentials at the subepicardium produced T-wave inversion(22). R1309 of Na_v_1.5 protein located in the S4 helices of DIII of α subunit of the sodium channel served as a voltage sensor. The S4 in all four domains involved in pore gating with different extents. The S4 helices in DI-DIII seem to influence channel activation, while the S4 of DIV seem to have a larger effect on inactivation than activation. The charged residues of S4 caused an outward motion of the helices, toward the extracellular side of the membrane, which in turn induced conformational changes within the pore. The S4 helices also had molecular interaction with surrounding S1-S3 helices, and thus facilitated activation(23). We hypothesized that the spatial and conformational changes of S4 in amino acid sequence induced by the harmful SCN5A mutation, R1309H for example, possibly led to abnormal outward motion toward the extracellular side of the membrane, or abnormal molecular interaction, which thus complicated with adverse consequence of pore gating abnormality with characteristics of drastically decreased peak density of I_Na_, depolarized activation and hyperpolarized inactivation. The S4 helices of DIII and DIV were the major determinants of the voltage dependence of lidocaine affinity(24). Notably, similar to L1308F mutation reported before(25), R1309H conferred predominant sensitivity to sodium blocker of lidocaine, by which induced more decreased of I_Na_, depolarized activation and hyperpolarized inactivation. It was also suggesting that lidocaine was a potential proarrhythmic drug for patient carrying R1308F, R1309H or R965C-R1309H mutations, and S4 helix of DIII might serve as a potential and sensitive site for targeting therapy.

However, in fact, it was most important and interesting for us, why and how the mutation of R1309H coexisting with R965C aggravated more deteriorative changes of pore gating property. Actually, a short amphiphilic α helix region (ranged from R954 to D979) in the DII/DIII linker resembling an S4 voltage sensor was an important determinant of sodium pore gating. The motif initiated from R961 to D979 (amino acid sequence of LALARIQRGLRFVKR) functioned as S4-like region carrying regularly arranged positive charges, including R965, R968, R971, and K974. Also, the potential phosphorylation site for PKA, an amino acid sequence of KRTT initiating from K974 to T977 site of Na_v_1.5 protein was located at the C-terminal end of this putative amphiphilic helix. This motif specifically affects the voltage dependence of current density, steady-state activation and inactivation of sodium channel after its deleting mutation (26). The substitution of R965C from the positively charged arginine to a neutral cysteine might influence channel gating properties, characteristics of hyperpolarized inactivation and altered recovery from inactivation due to impaired slow inactivation. Moreover, the three simultaneous substitutions of R965A, R968A, and R971A presented with predominantly decreasing of peak density of I_Na_(26). Furthermore, no matter the single mutation of R965C or R1309H, but also the compound mutations of R965C-R1309H, the Na_v_1.5 protein were obviously trafficking and locating to membrane expression. Based on the studies and phenomenon above, the compound mutations of R965C-R1309H synergistically influenced the pore gating properties of the sodium channel, suggesting that the important interaction between S4-like region in DII/DIII linker and S4 helix of DIII. Otherwise, although we hadn’t evaluated the effect of R965C-R1309H on the I_NaL_, the aggravating results were sufficient to explain the etiology for the complex clinical phenotypes in this family.

### Phenotypes and prognosis analysis in this family

It was worth noting that why the proband of II:3 suffered from recurrent ischemic strokes, even with an effective Warfarin therapy of INR2.0-3.0. Atrial standstill and ventricular pacing dependence were identified for II:3 during follow-up. Additionally, loss-of-function SCN5A mutations were commonly associated with ventricular dilatation and impairment in contractile function as reported before(27). We proposed that complete atrial dysfunction demonstrated by atrial standstill, potential ventricular contractile dysfunction, and mitral-tricuspid regurgitation caused by persistent ventricular pacing dependence were more likely to lead to thrombosis, which thus eventually increased the risk of recurrent cerebral embolisms. Consistent with this hypothesis, a study revealed that even though CHA_2_DS_2_-Vasc score was zero, two unrelated boys of 11 and 14-year-old carrying D1275N and E1053K mutations, respectively, had the occurrence of severe stroke related to atrial standstill due to potential and underlying atrial myopathy with characteristic of atrial fibrosis and impossible electrical capture of atrial muscle(28). Whether atrial standstill, complete atrial dysfunction or severe loss of function of sodium channel induced by single or multiple SCN5A mutations would increase the risk of recurrent embolism needs further clinical evaluation. III:7 had early onset of sick sinus syndrome, progressive atrioventricular block, and atrial fibrillation since 10-year-old. The carriers with Brugada syndrome associated with SCN5A mutation indicated the increased risk of progressive conduction disease, due to a similar and common mechanism of loss-of-function SCN5A mutation(29). III:3 at the age of 17 elicited apparent sinus bradycardia in the daytime with averaged sinus rate at less than 40 beats per minute, but without clinical symptoms so far, suggesting early sinus node dysfunction had occurred. Meanwhile, consistent with these phenomenons, familial cardiac conduction diseases caused by SCN5A mutation had profound male predominance and earlier age of onset(30).

In this family, the clinical phenotypes of carriers with heterozygous and compound mutations of R965C-R1309H were diverse and incomplete penetrance. II:2 was revealed with Brugada syndrome-like ECG, and occurred with syncope associated with overexertion and lack of sleep, while the other carriers had normal or slightly abnormal ECGs. This phenomenon was explained by several possibilities. For example, the compound mutations had an individual difference in the penetrance rate and genetic haploinsufficiency, that is, sodium current produced solely from the unaffected allele. The expression level of Na_v_1.5 protein and post-transcriptional modification in mutated and unaffected allele, cardiac structural remodeling induced by the fibrotic process, also some environmental factor (such as temperature and PH value) would influence the clinical phenotype and penetrance(31-33).

### Phenotypes associated with compound mutations of the SCN5A gene in previous studies

There were several limited types of research about the interaction of compound mutations. The T1620M or R1232W mutation only had a slight effect on sodium channel characteristics. Surprisingly, both coexisting mutations induced the disrupted membrane trafficking of Na_v_1.5 protein that was stuck in the endoplasmic reticulum, which thus resulted in non-functional sodium channel and eventually, Brugada syndrome occurred(34). Even though the W156X and R225W mutations were characterized as loss of function, respectively, the carriers with one of the two had no obvious clinical phenotype. Interestingly, when the compound heterozygous W156X and R225W mutations coexist, they induced more severe dysfunctional sodium channel, therefore the grievous clinical phenotypes including severe conduction disease, dilated cardiomyopathy and sudden death in young adults(35). The R34fs/60 mutation induced no sodium current, while R1195H had no effect on I_Na_ and I_NaL_, but abnormal voltage-dependent gating parameters. Both mutations coexisting surprisingly and significantly increased the I_NaL_, which may be the reason for QT prolongation and fever-induced ventricular arrhythmias for a 22-month-old boy(36). P1332L channel causing QT syndrome displayed hyperpolarized and slower inactivation, prominent I_NaL_, greater sensitivity to block by mexiletine. While A647D causing Brugada syndrome had no effect on the properties of I_Na_, it reduced I_Na_, I_NaL_ and the sensitivity to block by mexiletine, and even shorten the action potential duration when coexpressed with P1332L, which therefore induced different phenotypes of varying P wave/QRS aberrancy and mild QTc prolongation(37). P336L reduced I_Na_ by 85% of wild type, whereas I1660V abolished it. Only the proband carrying both mutations as compound heterozygosity exhibited Brugada syndrome. The I1660V channel trapped in intracellular organelles, likely in the endoplasmic reticulum, and could not properly traffic to the plasma membrane(38). Additionally, the two kinds of double mutations, including A735V and D1792N(39), P1048SfsX97 and T220I(40), could lead to sinus node dysfunction, sick sinus syndrome, atrial standstill and Purkinje fiber system disease, but the interaction and electrophysiological mechanism in double mutations were unknown. Interestingly, not all double mutations or multiple mutations would aggravate the clinical phenotype. For example, H558R was a common polymorphism in lone atrial fibrillation, also as a disease-modifying mutation, and eliminated the harmful effect of P2006A mutation as a gain of function through weakens the I_NaL_. In the other side, it would recover the function of sodium channel of R282H mutation as loss of function, through enhancing peak current of I_Na_. In another study, H558R could recover the membrane expression of Na_v_1.5 and I_Na_ in the cells of D1690N mutation, while had no obvious recovery on I_Na_ in the cells of G1748D mutation(41). In addition, not all the compound mutations showed obvious clinical phenotype until inappropriate drug therapy, especially the antiarrhythmic drug. The compound mutations of V232I-L1308F produced no changes of sodium channel properties, consistent with lack of phenotype. The lidocaine application made the sodium channel containing V232I-L1308F mutation change to decreased peak current of I_Na_, more depolarized activation and hyperpolarized inactivation, which therefore induced Brugada syndrome. The underlying mechanism was that V232I and L1308F located in S4 of DI and DIII, respectively, and conferred great sensitivity of sodium channel to lidocaine through altering the coupling between S4 and S6 to increase the affinity for lidocaine. However, it remains unknown about the underlying mechanisms by which how the mutations interacted with each other and influenced the properties of the sodium channel and how the intracellular alteration and motion of pore gating change.

## Materials and Methods

### Patient phenotyping

This study was approved by the Guangdong Medical Institutional Review Board and Medical Ethics Committees [No.GDREC2016001H (R1)]. Informed written consent was obtained from all patients. Baseline 12-lead electrocardiographs (ECGs) and ambulatory 24-hour Holter ECGs were recorded at a paper speed of 25 mm/s in all subjects. According to the current guidelines, Brugada syndrome was defined as an ST-segment elevation with type 1 morphology for ECGs [2 mm in, at least, one lead among the right precordial leads V_1_ and V_2_, positioned in the second, third or fourth intercostal space]. Sinus node disease was defined in one of the following cases: sinus bradycardia <40 beats/min while awake, sinoatrial block or sinus arrest with pauses >3.0 seconds, bradycardia-tachycardia syndrome, defined as episodes of atrial tachyarrhythmia coexisting with sinus bradycardia, sinoatrial block, or sinus arrest, and symptoms of syncope or dizziness.

### Genetic analysis

Genomic DNA was extracted from peripheral blood leukocytes using standard protocols. The coding and splice-site regions of genes susceptible to hereditary arrhythmia and cardiomyopathies(10) were sequenced for probands II:3 and III-7 using Illumina HiSeq2500 Analyzer (Illumina, San Diego, CA, USA)(11), and assessed with Burrows-Wheeler Aligner(12), SOAPsnp and SAMtools software(13). DNA variants were considered disease-causing rather than polymorphisms if: (1) absent from the 1,000 Genomes exome database (http://www.1000genomes.org), (2) <1% in the Exome Variant Server (EVS) of the NHLBI Exome Sequencing Project, which includes patients with cardiac diseases [http://evs.gs.washington.edu/EVS/ (02-2013 accessed)] and (3) present in highly conserved regions (http://genome.cse.ucsc.edu/index.html)(11). Gene mutations were annotated based on the following cDNA reference sequences from Ensemble: SCN5A NM_001099404.1.

### Stable cell line construction

SCN5A plasmids for the mutant sodium channels (p.R1309H/p.R965C/p.R1309H-p.R965C) were first using dsDNA oligo (GEN CHEM, Shanghai) and used as the template in the subsequent polymerase chain reaction (PCR) synthesized using 2μl 10×buffer, 2.5mM dNTPs, 0.4μl Primer(+), 0.4μl Primer(-), 0.2 μl Taq polymerase, 1μl Template and 15.2μl ddH_2_O. The PCR product for SCN5A was first cloned into Ubi-MCS-3FLAG-SV40-EGFP-IRES-puromycin (GEN CHEM, Shanghai) with Age I/EcoR I and subcloned into the BgI II recognition with a sequence of pBudCE4.1 (Invitrogen) with BamH I/BgI II. The subclone procedure allowed bicistronic expression of Enhanced Green Fluorescent Protein (EGFP) and puromycin resistant gene under the control of EF-1α promoter. The PCR product of SCN5A was cloned into the pBudCE4.1 with Hind III/Xba I under the control of Ubi promoter. Besides, three flag epitopes DYKDDK was introduced in the frame at the C-terminus of SCN5A. The base sequence of the SCN5A clone was analyzed and identical to the published SCN5A (hH1, NM_198056) which were detected by PCR procedures.

### Electrophysiological recordings

HEK293T cells were recorded with an Axopatch 700B amplifier controlled by pClamp 10.6 and acquisition frequency of the samples at 10 kHz. Experiments are performed at the temperature of 32°C with TC-324B thermostat cell trough. The internal solution included (in mM): CsCl 50, CsF 30, L-aspartic acid 50, NaCl 10, tetraethylammonium hydroxide (TEA-OH) 11, EGTA 5, HEPES 10, PH 7.3 adjusted with CsOH. And the external solution contained (in mM): NaCl 120, KCl 5.4, CaCl_2_ 1.8, MgCl_2_ 1, HEPES 10, glucose 10, tetraethylammonium chloride (TEA-Cl) 20, PH adjusted with NaOH. Pipettes were made from borosilicate glass through Sutter P-97 Micropipette Puller, with the pipette resistance ranged 2.5MΩ to 5.0MΩ when filled with an internal solution. And series resistance was less than 20MΩ.

In order to detect the activation characteristics of sodium current, we used a single pulse voltage stimulation method: sodium current was induced by a 50 ms depolarization stimulation, which ranged from −100mV to +60mV with the increment of 5mV at the holding potential of −120mV. As for the steady-state inactivation characteristics of sodium current, a two-pulse protocol was performed: the first pulse with the duration of 500ms from −140mV to −40mV at 5mV increment followed in succession by the second pulse fixed at −40mV, and the sodium current induced by the second pulse was the targeted current. Both steady-state activation and inactivation curves were fitted with Boltzmann function: I/Imax=(1+exp(-(V-V_1/2_)/k)) ^−1^ and I/Imax=(1+exp((V-V_1/2_)/k)) ^−1^ respectively, I/Imax is the induced currents normalized to its maximal value, V is the test potential, V_1/2_ is the test voltage at which the current amplitude is the half-maximal, and K is the slope factor. Time constant tau (τ) for steady-state activation and inactivation both obtained from the Clampfit software.

### Localization of sodium channels

To identify trafficking components, which may be caused by double-mutant Na_v_1.5 channels, we used confocal microscopy to localize Na_v_1.5 channels conjugated to a green fluorescent protein (GFP). HEK293T cells were cultivated on 35-mm cell culture dishes and forty-eight hours after cell thawing, the culture medium was removed and cells were washed with ice-cold PBS. The cells were then fixed by incubation in 4% paraformaldehyde at room temperature for 30 min and permeabilized with PBS containing 0.5% Tween 20 and 10% Normal goat serum for another 30 min. Before confocal imaging, cells were incubated in PBS containing primary antibody (anti-flag 1:500, CST) overnight at 4°C and secondary antibody goat anti-Rabbit IgG conjugated with Alexa Fluor 594 (1:2000, CST) for 30 min. Confocal images were obtained and analyzed a Zeiss LSM 800 with Airyscan confocal microscope (Zeiss, Jena, Germany) and images were acquired with a FluoView acquisition software program on a microcomputer.

### Statistical analysis

Data analysis was performed with the corporation of Clampfit 10.6 and GraphPad Prism 5. All the data were presented as mean ± SEM. For comparisons, both slope factor (k) and V_1/2_ were calculated using one way ANOVA followed homogeneity test for variance, while the rest of the data were calculated using the unpaired two-sample Student’s T-test. *A p*-value less than 0.05 were considered to be statistically significant.

## Acknowledgments

We are grateful to BGI (Shenzhen, China) for technical assistance.

